# Measurement of the Glutathione Redox Potentials in Membraneless Organelles

**DOI:** 10.1101/2025.03.30.646167

**Authors:** Anna Rovira, Pablo Rivera-Fuentes

## Abstract

Glutathione (GSH) is essential for cellular redox regulation; however, the redox potential (E_GSH_) of the nucleus and membraneless organelles (MLOs) remains poorly understood. Here, we use the Grx1–roGFP2 sensor to measure the E_GSH_ of the nucleus and several MLOs such as the nucleoli, stress granules, p-bodies, paraspeckles, and Cajal bodies. Unlike suggested by previous findings, we found that nuclear E_GSH_ is stable throughout the cell cycle and identical to the cytosolic E_GSH_, and the E_GSH_ of MLOs mirrors their surrounding environment. These findings challenge existing paradigms and provide novel insights into redox homeostasis across subcellular compartments.

## Introduction

Glutathione (GSH) plays a central role in maintaining cellular homeostasis and redox balance.^[1]^ GSH is a tripeptide composed of glutamate, cysteine, and glycine and is the most abundant low-molecular-weight, thiol-containing metabolite in most organisms,.^[2]^ It functions primarily as an antioxidant, with intracellular concentrations reaching up to 20 mM in cells.^[3]^ Within the cell, glutathione exists almost exclusively (99%) in its reduced form (GSH), while only 1% is found as oxidized glutathione (GSSG). The dynamic balance between GSH and GSSG, often expressed as the glutathione redox potential (E_GSH_), is a critical indicator of the oxidative state of the cell.^[4–6]^ Beyond its antioxidant properties, GSH contributes to various cellular processes, including DNA and protein synthesis, cell proliferation, apoptosis, and signal transduction.^[7]^

The synthesis of GSH occurs exclusively in the cytosol, from where it is transported to different subcellular compartments, such as the endoplasmic reticulum (ER), Golgi apparatus, mitochondria and the nuclear matrix.^[8–10]^ Notably, the concentration of GSH and the GSH/GSSG ratio vary across these organelles, reflecting compartment-specific redox environments that are tightly regulated by the cell.^[11]^ For instance, the cytosol typically exhibits a E_GSH_ between -315 and -325 mV,^[12,13]^ with GSH concentrations ranging between 1 and 20 mM,^[3]^ whereas mitochondria possess an independent redox pool characterized by a narrow GSH concentration range (5-10 mM), and a significantly lower redox potential of approximately -360 mV.^[14]^ In contrast, organelles of the secretory pathway are more oxidizing, with the ER displaying a E_GSH_ = -208 mV^[15]^ and the Golgi apparatus a E_GSH_ = -157 mV.^[16]^

Current figures for intra-organelle GSH redox potentials have been measured using genetically encoded fluorescent sensors based on redox-sensitive green fluorescent proteins (roGFPs). These tools have revolutionized our ability to monitor subcellular redox dynamics in real time.^[15,17]^ roGFPs are engineered to have two cysteine residues whose oxidation to form a disulfide induces structural changes that alter their fluorescence properties. The oxidation and reduction of roGFPs is driven by small-molecule thiols/disulfides such as the GSH/GSSG pair, thus enabling the quantification of their redox potential through the Nernst equation.^[18,19]^ Coupling roGFPs to glutaredoxin (Grx) enhances the sensitivity and selectivity towards GSH/GSSG, enabling the organelle-specific measurement of E_GSH_.^[12,14]^ Thus, these sensors are uniquely positioned to provide insights into the redox states of organelles, study the complexity of compartmentalized redox regulation, and either confirm or challenge long-standing paradigms in redox cell biology.^[20]^

Grx1-roGFP sensors have been used to measure characterize the GSH redox potential in various membrane-bound organelles.^[16,20]^ In contrast, the redox characteristics of membraneless organelles (MLOs)^[21]^ remain largely unexplored. MLOs have gained prominence recently because of their ability to concentrate specific proteins, nucleic acids, and small molecules in specific subcellular locations without the need for a lipid membrane.^[22]^ Owing to their dynamic formation, they can respond rapidly to changes in the cellular environment, including temperature, pH, or post-translational modifications.^[23]^ MLOs play a key role in gene expression and RNA processing,^[24]^ and their dysregulation has been linked to amyotrophic lateral sclerosis,^[25]^ Alzheimer’s and Parkinson’s diseases,^[26,27]^ among other diseases.^[28]^ Recent studies have reported the accumulation of specific small molecules in membraneless organelles,^[29]^ and it seems like the properties of the small molecules define the extent to which they partition into biomolecular condensates.^[30]^ Thus, a key question in redox biology is whether the two components of a redox buffer (e.g., GSH and GSSG) partition similarly or not into MLOs. If there is no substantial difference between their partition coefficients, the redox environment of the MLO would be similar to its immediate environment (e.g., cytosol or nucleoplasm). However, if one component of the redox buffer partitions selectively into the MLO, that would lead to a very different redox environment in the MLO compared to its environment. In this study, we set out to answer this question using a redesigned Grx1–roGFP2 sensor that can be robustly targeted to all these MLOs.

## Results and Discussion

### Engineering and Validation of the miniGrx1-roGFP2 Biosensor

In our hands, the original Grx1-roGFP2 biosensor displayed somewhat inconsistent performance when fused to various targeting sequences for subcellular localization. Some of the problems that we encountered (data not shown) were dim signals, inaccurate localization, and low expression levels. We hypothesized that the properties of this fusion protein were negatively impacted by the long, unstructured linker (six consecutive GGGS units) between the Grx1 and roGFP2 proteins. This flexible linker, however, was deemed important for the functionality of the sensor because it allowed the two proteins to find an optimal orientation for G1×1 to glutathionylate roGFP2.^[13]^ We posited that we could design a structured linker that aligned the two proteins in an optimal conformation, preserving the functionality of the original sensor while creating a more compact fusion with better expression and targetability.

We started by predicting the structure of the original sensor using AlphaFold2 (AF2)^[31]^ (Figure 1a). This prediction confirmed the unstructured nature of the original linker. Computationally, we removed this linker and manually reorient Grx1 so that GSSG binding site was close and aligned with the two reactive cysteines of roGFP2. Using this re-oriented structure, we employed RosettaFold Diffusion (RFD)^[32]^ to inpaint a new linkers between the two proteins. RFD generated backbone coordinates for a few linkers with lengths between 20 and 30 amino acids. We then used ProteinMPNN^[33]^ to predict which sequences of amino acids would preferably fold into the backbone geometries designed with RFD. Finally, with the new sequences provided by ProteinMPNN, we calculated the structure of the new sequences using AF2. We ranked the models by their predicted local distance different test (pLDDT) and selected the top two models of different linker lengths (20 and 30 amino acids) to test experimentally.

**Figure 1.**
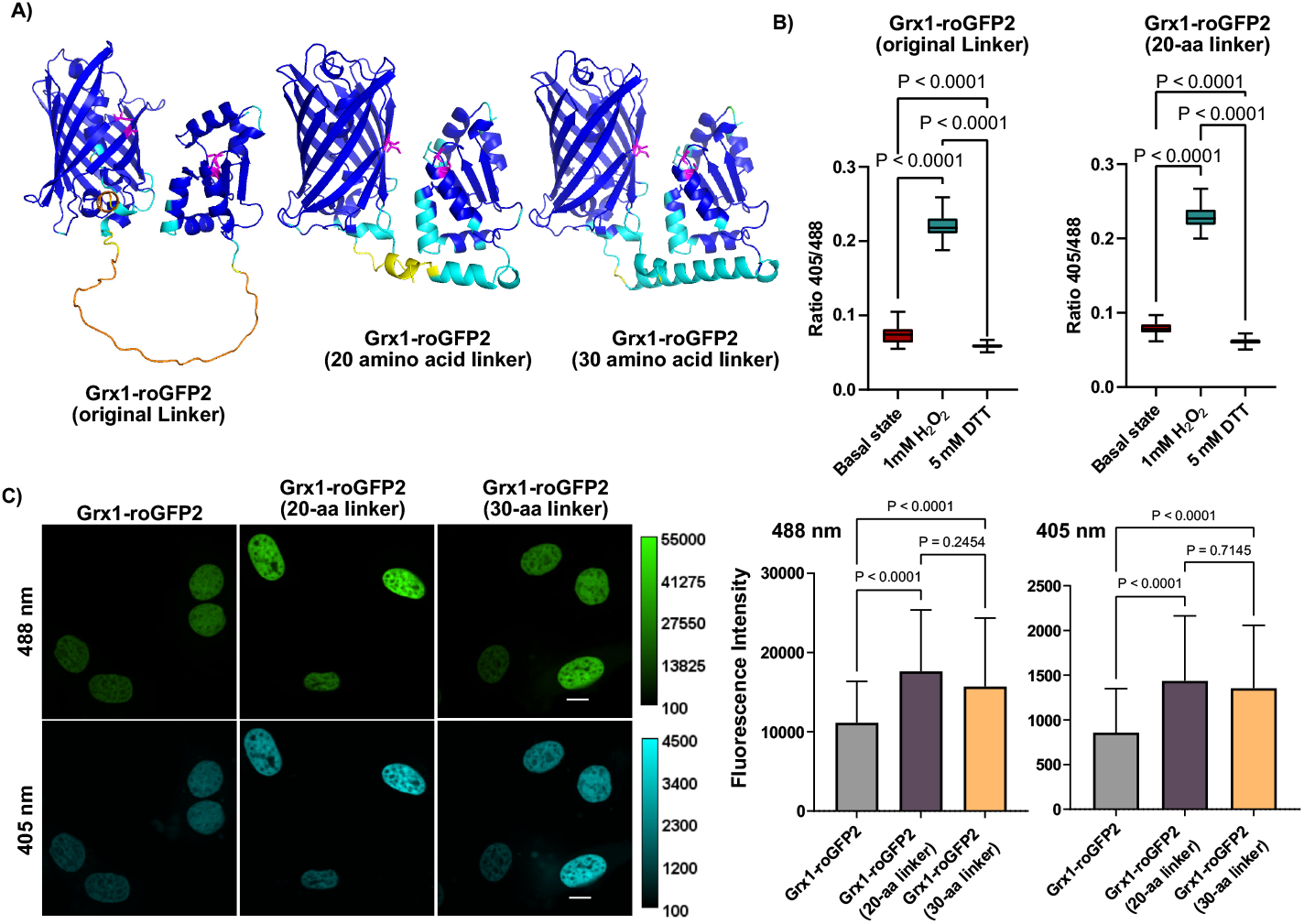
Engineering and Validation of the Redesigned Grx1-roGFP2 Biosensors. **(A)** AF2 structural models comparing the original Grx1-roGFP2 sensor (left) to the redesigned versions featuring 20- and 30-amino-acid linkers (middle and right, respectively). The models are color-coded by AF2’s per-residue pLDDT, with blue indicating regions of higher confidence and red indicating lower confidence. The magenta residues highlight the reactive cysteines in roGFP2 and Grx1. **(B)** Functional redox responsiveness of the original and the 20-amino-acid linker redesigned sensor, assessed by the fluorescence ratio (405 nm / 488 nm) under basal (untreated), oxidizing (1 mM H_2_O_2_), and reducing (5 mM DTT) conditions. Tukey-style box plots are presented as boxes that span from the 25th to the 75th percentiles with the median marked by a horizontal line, while whiskers extend to the most extreme observations within 1.5 times the interquartile range, and values outside this range are identified as outliers. Statistical significance was determined using one-way ANOVA (Tukey’s multiple comparisons test) with N > 65 independent cells examined (**Table S5**). The box plots show that all variants exhibit reversible shifts in the fluorescence ratio, confirming that the redesigned Grx1-roGFP2 constructs retain robust redox sensitivity comparable to the original. **(C)** Comparison of fluorescence intensity and localization between the original and redesigned sensors in HeLa cells. **Left**: Representative fluorescent microscopy images acquired at excitation wavelengths of 405 nm (pseudo-colored in cyan) and 488 nm (pseudo-colored in green) with emission collected at 525/50 nm. Images were taken under basal conditions and 3 min after treatment with 1 mM H_2_O_2_ (oxidizing conditions) or 5 mM DTT (reducing conditions). The redesigned sensors show both enhanced expression and accurate nuclear localization similar to the original construct. Scale bar: 10 μm. **Right**: Quantitative bar charts illustrating the mean fluorescence intensities (± SD) for each sensor in the 405 nm and 488 nm channels. The data demonstrate a significant increase in fluorescence intensity for the redesigned constructs compared to the original, and not statistically significant difference among both redesigned constructs.

We cloned the original sensor and the two computational designs into vectors containing a fragment of histone H2B for nuclear targeting in mammalian cells (Figure 1c). A simple assessment of brightness in cells revealed that both redesigned sensors showed a significant increase in fluorescence intensity across both excitation channels, consistent with enhanced protein expression (**Figure 1c** and **Figure S3**).

We validated the functionality of the new biosensors by exposing transfected cells to controlled redox challenges. Oxidizing conditions were induced by exposing the cells to hydrogen peroxide (1 mM H2O2), while we exposed cells to dithiothreitol (5 mM DTT) to induce reducing conditions. Both the parent and redesigned sensors responded as expected, with the new sensors demonstrating intact functionality, as evidenced by reversible changes in fluorescence corresponding to redox potential shifts (**Figure 1b** and **Figure S2**).

Importantly, localization studies revealed that the modified biosensors maintained accurate targeting to their respective organelles. For further validation, the redesigned sensors were also targeted to the cytosol and the Golgi apparatus, which are compartments for which E_GSH_ values are well-documented in the literature (**Figure 2** and **Figure S1**).^[16]^ Overall, we find that the measured redox potentials in these compartments were in good agreement with published data, confirming the reliability and versatility of the redesigned sensors. Collectively, these results establish the superiority of the redesigned Grx1-roGFP2 biosensors in terms of expression, fluorescence performance, and functional integrity.

**Figure 2.**
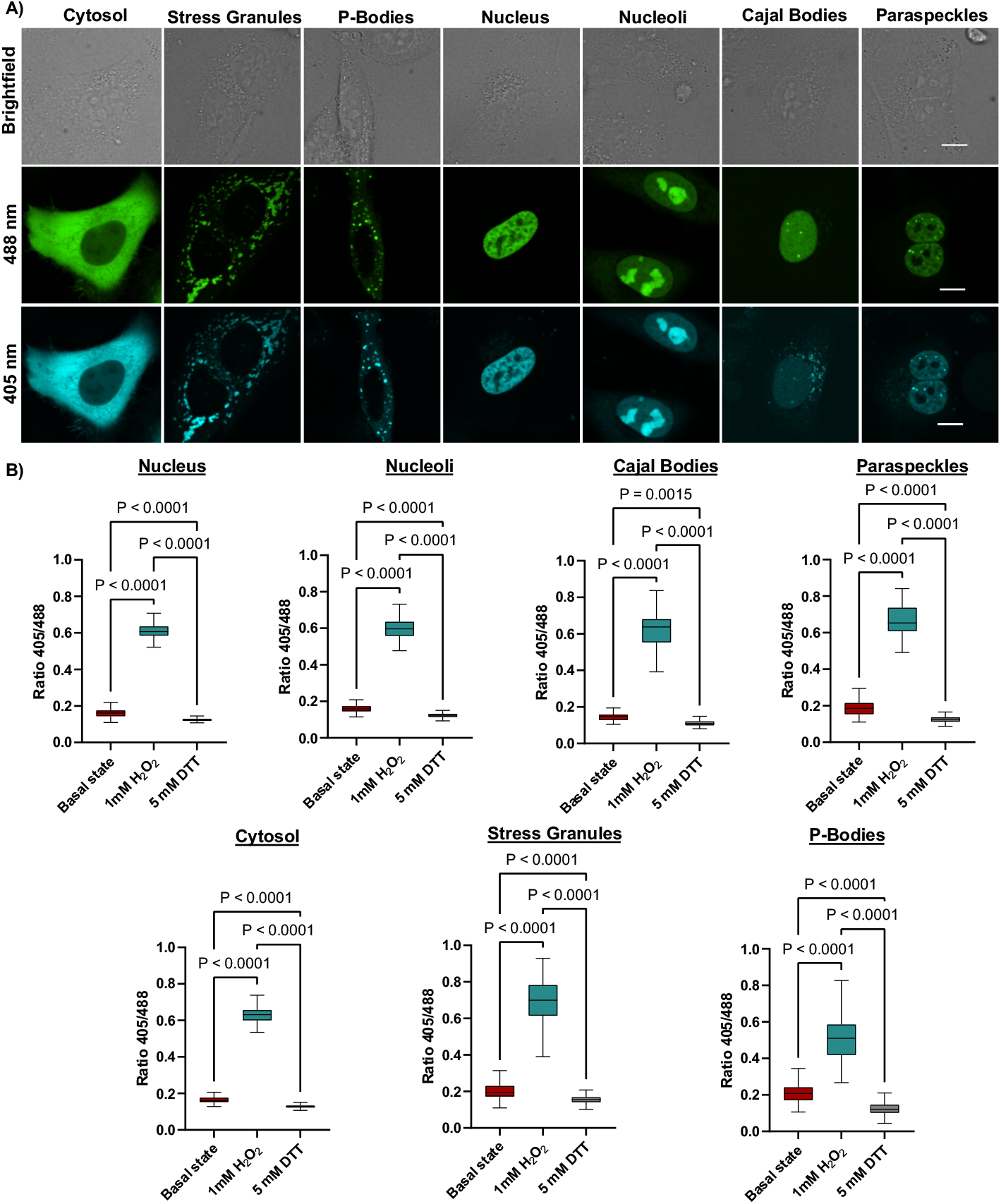
Localization and Functional Assessment of the 20-Amino-Acid Redesigned Grx1-roGFP2 Biosensor in HeLa Cells. **(A)** Confocal images showing the targeted expression of Grx1**-** roGFP2 biosensor in distinct compartments of HeLa cells. From left to right, the images display expression in the cytosol, stress granules, processing bodies, nucleus, nucleoli, Cajal bodies, and paraspeckles. For each compartment, the bright field image is shown at the top, followed by the 488 nm channel (pseudo-colored in green) and the 405 nm channel (pseudo-colored in cyan) at the bottom with emission collected at 525/50 nm. Scale bar: 10 μm. **(B)** Functional redox responsiveness of the sensor was assessed by the fluorescence ratio (405 nm/488 nm) under basal (untreated), oxidizing (3 min, 1 mM H_2_O_2_), and reducing (3 min, 5 mM DTT) conditions. Tukey-style box plots are presented as boxes that span from the 25th to the 75th percentiles with the median marked by a horizontal line, while whiskers extend to the most extreme observations within 1.5 times the interquartile range; values outside this range are identified as outliers. Statistical significance was determined using one-way ANOVA (Tukey’s multiple comparisons test) with N > 65 independent cells examined (**Table S6**). For the nucleus (N > 80 cells) and cytosol (N > 139 cells), N indicates the number of cells examined; for nucleoli (N > 214), Cajal bodies (N > 56), paraspeckles (N > 82), stress granules (N > 462), and processing bodies (N > 521), N corresponds to the number of individual membraneless organelles analyzed. (This is replicate 1; replicates 2 and 3 are provided in the Supplementary Information.)

### Measurement of Glutathione Redox Potential in Membraneless Organelles and Their Surroundings

Given the superior performance and smaller size of the redesigned sensor with a 20-amino-acid linker (Figure 1), we used it for targeting MLOs and called is miniGrx1-roGFP2. This biosensor was targeted to specific subcellular regions, including the nucleoli, stress granules, p-bodies, nuclear paraspeckles and Cajal bodies, along with their respective surrounding environments, i.e., the cytosol and nucleoplasm. Targeting was achieved by fusing the biosensor to well-established localization signals. For instance, a nucleolar localization signal (NoLS) was used for nucleoli,^[34,35]^ while protein markers such as G3BP1, Dcp1b, Coilin, and PSPC1 were employed for stress granules,^[36]^ p-bodies,^[37]^ Cajal bodies, and paraspeckles,^[38,39]^ respectively. For nuclear and cytosolic targeting, we used histone H2B and a nuclear export sequence (NES),^[3,40]^ respectively.

The redox functionality of miniGrx1–roGFP2 in each compartment was verified in HeLa cells transfected with the corresponding plasmids. Imaging was performed 24 hours post-transfection using excitation wavelengths of 405 nm and 488 nm, allowing calculation of the redox state under basal conditions, after oxidation with hydrogen peroxide (1 mM H2O2), and following reduction with dithiothreitol (5 mM DTT). The fluorescence intensity ratios (405 nm/488 nm) obtained under these conditions were used to calculate the degree of oxidation of roGFP2 (OxD_roGFP2_) and subsequently convert it to E_GSH_ using the Nernst equation (Eq. 1).^[15]^

The redox potential (E_GSH_) was calculated using the following equation:

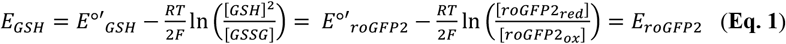

Where *E*°’(roGFP2) is the standard redox potential of the fluorescent protein (−280 mV, at pH 7),^[12]^ *R* is the gas constant (8.315 J K^-1^ mol^-1^), *T* is the temperature (K), and *F* is the Faraday constant (96’485 C mol^-1^).^[41]^ The oxidation status of the glutathione pool is derived from the OxD_roGFP2_ value, calculated as (**Eq. 2**):

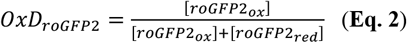

Experimentally OxD_roGFP2_ is directly calculated from the fluorescence intensity ratio between the 405 nm and 488 nm excitation channels, using the following equation (**Eq. 3**):

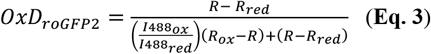

Here, R_red_ and R_ox_ are the fluorescence ratios under fully reduced (DTT-treated) and fully oxidized (H2O2-treated) conditions, respectively, while I_488red_ and I_488ox_ are the fluorescence intensities at 488 nm in reduced and oxidized states. Once OxD_roGFP2_ is determined, it is used in the Nernst equation to compute E_GSH_:

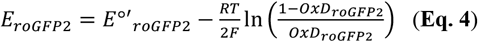

This approach ensures precise quantification of E_GSH_ values in different compartments under varying conditions. miniGrx1-roGFP demonstrated reversible fluorescence changes in response to oxidizing and reducing treatments, confirming its redox activity across all MLOs and their surrounding compartments (**Figure 2b**). Fluorescence microscopy also confirmed that miniGrx1-roGFP2 localized appropriately to their respective targets and maintained functionality throughout the experiments (**Figure 2a**).

The calculated E_GSH_ values for the nucleoli, stress granules, p-bodies, Cajal bodies, and paraspeckles, as well as their surrounding compartments, are summarized in **Table 1**, with representative fluorescence intensity data and imaging results shown in **Figure 2a**.

**Table 1.** E_GSH_ Values Across Subcellular Compartments and MLOs. E_GSH_ values (in mV) are shown for stress granules, processing bodies, nucleoli, nuclear paraspeckles, and Cajal bodies, along with the corresponding surrounding compartments (cytosol for stress granules and processing bodies; nucleus for nucleoli, paraspeckles and Cajal bodies). Data were calculated using the Nernst equation. The standard deviations (SD) represent uncertainties derived from propagating the experimental errors through the Nernst equation.

| Region | $E_{\text{GSH}}$ (mV) | Surrounding Compartment | $E_{\text{GSH}}$ (mV) |
| --- | --- | --- | --- |
| Stress Granules | $-295 \pm 22$ | Cytosol | $-300 \pm 13$ |
| Processing Bodies | $-287 \pm 22$ | | |
| Nucleoli | $-300 \pm 19$ | Nucleus | $-301 \pm 22$ |
| Paraspeckles | $-294 \pm 20$ | | |
| Cajal Bodies | $-306 \pm 23$ | | |

The E_GSH_ measurements revealed striking uniformity between the redox potentials of MLOs and their surrounding compartments. For instance, nucleoli within the nucleus displayed an E_GSH_ of -300 ± 19 mV, closely matching the nuclear E_GSH_ of -301 ± 22 mV. Similarly, stress granules and p-bodies exhibited E_GSH_ values nearly identical to the cytosolic redox potential. These findings strongly suggest that MLOs do not establish distinct redox microenvironments but rather equilibrate with the surrounding redox conditions. The absence of redox compartmentalization in MLOs may be explained by their biophysical nature; as liquid-liquid phase-separated domains,^[21]^ MLOs lack physical barriers like lipid membranes that could isolate them from the bulk cytosol or nucleoplasm. Additionally, GSH and GSSG only differ by the presence of a thiol or a disulfide and their molecular weight. Moreover, since differential partitioning of small molecules into MLOs is largely driven by solubility and hydrophobicity,^[30]^ there should not be a significant difference in how GSH and GSSG partition into MLOs, maintaining the equilibrium of the surrounding environment.

Overall, these findings provide new insights into redox homeostasis within eukaryotic cells, emphasizing the non-compartmentalized nature of redox regulation in membraneless organelles. From a functional perspective, the uniform redox potential between MLOs and their surroundings may facilitate seamless coordination of redox-dependent processes, such as RNA metabolism in stress granules or ribosome assembly in nucleoli. This homogeneity likely supports cellular resilience, ensuring that fluctuations in redox state are uniformly sensed and managed across compartments.

### Analysis of Glutathione Redox Potential in the Nucleus and Nucleoli During the Cell Cycle

Building on the observation that the E_GSH_ in the nucleus and nucleoli remains uniform and equilibrated with their surroundings under steady-state conditions, we extended our analysis to investigate potential fluctuations during the cell cycle. The dynamic processes of cell division, involving chromatin condensation and nuclear envelope breakdown, could influence redox homeostasis within these compartments. However, our findings indicate that both the nucleus and nucleoli maintain stable E_GSH_ values throughout the cell cycle, reinforcing the robustness of redox regulation in these regions.

Using Grx1-roGFP2 targeted to the nucleus, we performed a 48-hour timelapse imaging to track changes in the 405/488 fluorescence ratio, which correlates with E_GSH_ (**Figure 3a** and **Figure 3c**). Overall, the 405/488 ratio remains stable, with marked peaks that coincide with the onset of mitosis. This increase occurred as cells adopted a rounded morphology and detached from the surface, a hallmark of mitotic entry.^[42]^ However, this peak diminished once cell division was complete and the daughter cells reattached to the substrate. The perfect correlation between the ratio change and the detachment and reattachment of the cell to the coverslip made us suspect that what we were observing was an imaging artifact.

**Figure 3.**
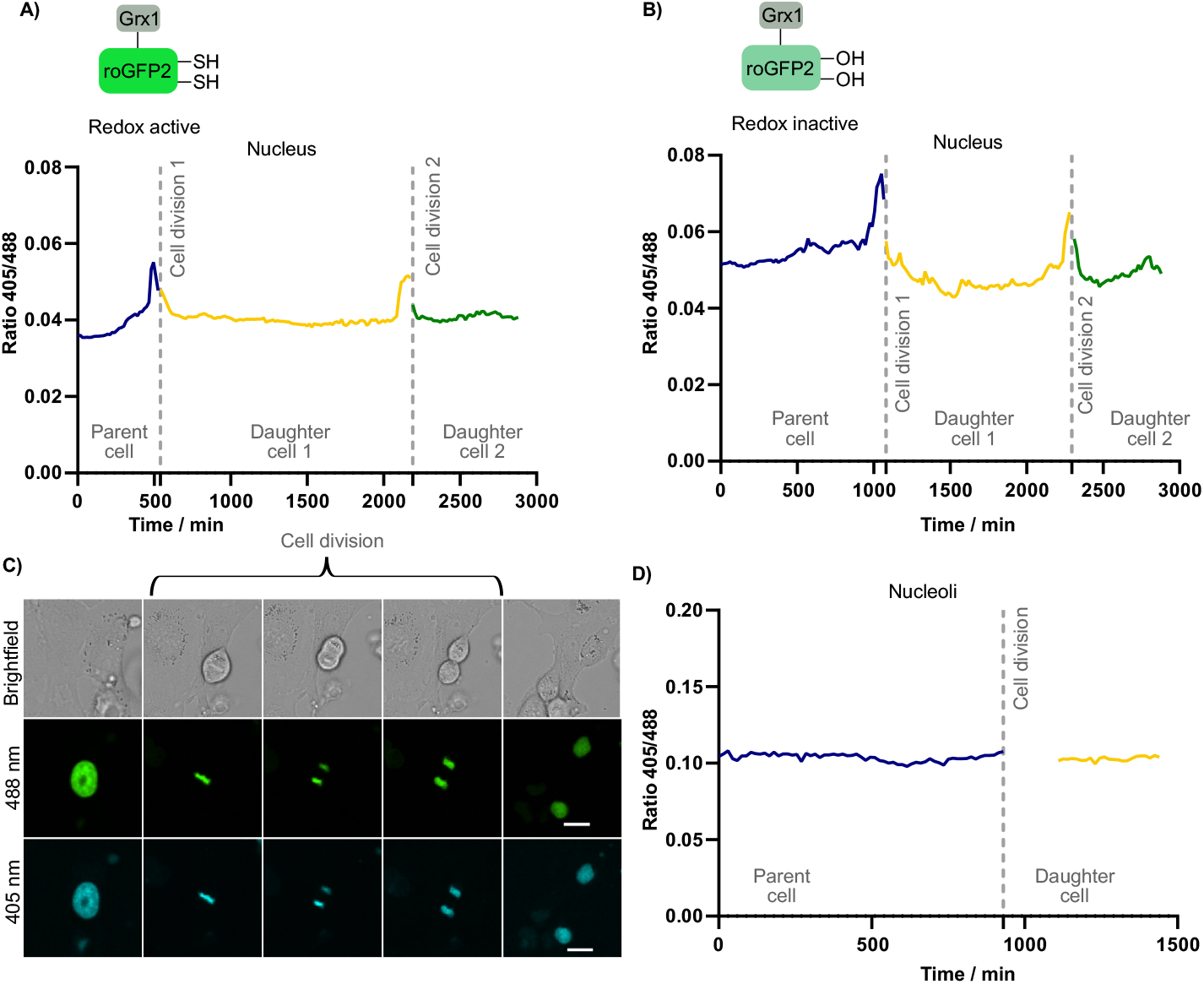
Dynamic Changes in Redox State During Cell Division. **(A)** Variations of the 405/488 fluorescence ratio in the nucleus over two cell division events monitored within 48 hours using the redox-active sensor. The original cell is depicted in blue, with the corresponding daughter cells post-division shown in yellow and green. **(B)** Similar analysis using the redox-inactive sensor, with the original cell in blue and the corresponding daughter cells in yellow and green. **(C)** Representative confocal images of the nucleus during cell division, acquired using the Grx1-roGFP2 sensor. Images include a brightfield view, and fluorescence channels at 405 nm and 488 nm with emission collected at 525/50 nm. Scale bar: 20 μm. **(D)** Variations of the 405/488 fluorescence ratio in the nucleoli during a single cell division event, with the original cell shown in blue and the selected daughter cell in yellow.

To determine whether this transient ratio change reflected genuine redox shifts or imaging artifacts, we introduced mutations to roGFP2, replacing its cysteine residues with serines, thus rendering it inactive. This inactive roGFP2 construct was expressed in HeLa cells and subjected to the same 48-hour timelapse imaging (**Figure 3b**). The resulting fluorescence patterns closely resembled those observed with the redox-sensitive H2B-Grx1-roGFP2 sensor, suggesting that the ratio increase during mitosis is not related to changes in E_GSH_ and thus, it likely resulted from focus adjustments due to cell detachment and rounding during division. As such, these results suggest that the redox potential in the nucleus remains stable across all phases of the cell cycle, irrespective of structural and morphological changes associated with mitosis.

A parallel series of experiments was conducted using the nucleolar-targeted Grx1-roGFP2 original biosensor. Timelapse imaging revealed no significant changes in the 405/488 fluorescence ratio in the nucleoli during the cell cycle (**Figure 3d**).

This stability in nucleolar E_GSH_ further corroborates the uniformity of redox regulation across MLOs and their surrounding compartments. Despite the dynamic reorganization of nucleolar components during mitosis, the redox potential appears tightly maintained, likely reflecting efficient equilibration with the nuclear and cytosolic environments. This stable E_GSH_ observed in the nucleus and nucleoli throughout the cell cycle has important implications for cellular redox homeostasis, as these compartments host critical processes such as DNA replication, transcription, and ribosome biogenesis, all of which require finely tuned redox regulation.^[24,43]^ Our findings suggest that the redox potential of the nucleus and nucleoli is tightly buffered, ensuring that redox-sensitive biochemical pathways operate consistently despite structural and morphological changes during mitosis.

Moreover, the absence of significant redox fluctuations does not necessarily disprove the hypothesis that cell cycle progression is accompanied by shifts in nuclear GSH concentration^[44,45]^ because the total concentration of GSH could change without altering its redox potential. Nevertheless, further studies will be necessary to test this hypothesis again in light of the results presented herein.

## Conclusions

This study presents the first comprehensive measurements of the E_GSH_ in MLOs and their surrounding compartments, providing novel insights into the mechanisms of redox homeostasis at the subcellular level. By utilizing an engineered Grx1–roGFP2 biosensor optimized for fluorescence intensity and targetability, we systematically quantified E_GSH_ in the nucleus, nucleoli, paraspeckles, Cajal bodies, stress granules, p-bodies, and the cytosol.

Our findings support the view that small molecules with similar solubilities and hydrophilicities partition similarly into MLOs, with the important consequence that in the case of the GSH/GSSG redox buffer, MLOs do not display specific redox environments and instead the redox properties of these organelles closely mirrors that of their surroundings. Notably, the nuclear E_GSH_, despite the nucleus being a membrane-bound compartment, is indistinguishable from that of the cytosol and remains remarkably stable throughout the cell cycle. Collectively, these results redefine our understanding of redox homeostasis across subcellular compartments by highlighting a consistent E_GSH_ that supports essential cellular processes such as DNA replication, transcription, and ribosome biogenesis.

## CRediT authorship contribution statement

**Anna Rovira:** Writing – review & editing, Writing – original draft, Validation, Methodology, Investigation, Formal analysis, Conceptualization. **Pablo Rivera-Fuentes:** Writing – review & editing, Writing – original draft, Validation, Methodology, Investigation, Formal analysis, Conceptualization.

### Notes

The authors declare no competing financial interest. All new plasmids generated in this work are available through Addgene: #236545-236551.

