## Supplementary Information for "Measurement of the Glutathione Redox Potentials in Membraneless Organelles"

#### 1. Supplementary Figures

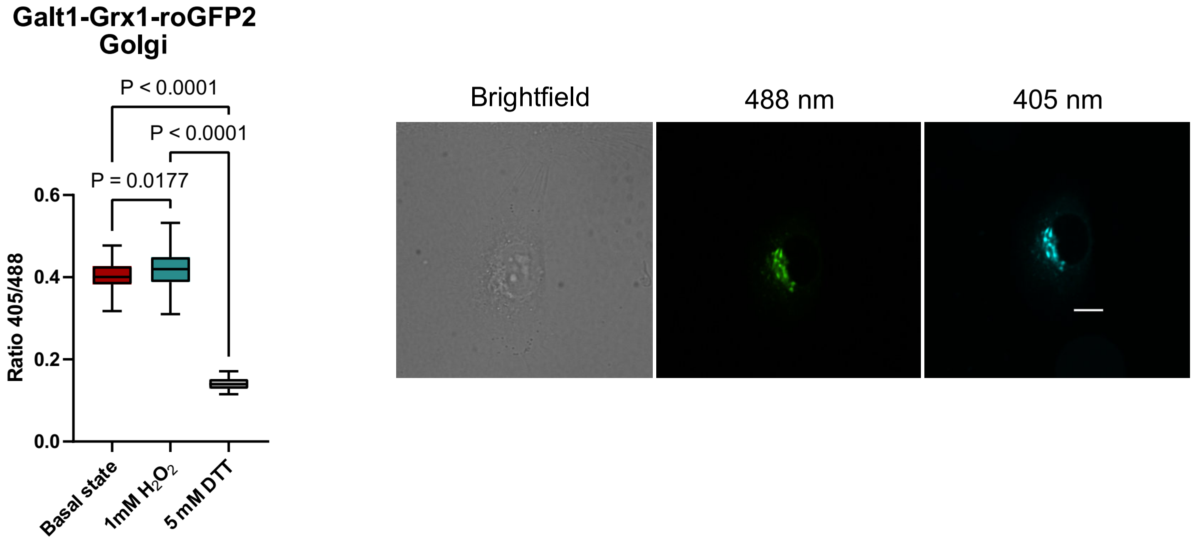


**Figure S1. Calculation of Golgi Redox Potential. Left**: Functional redox responsiveness of GALT1-miniGrx1-roGFP2 sensor was assessed by the fluorescence ratio (405 nm / 488 nm) under basal (untreated), oxidizing (3 min, 1 mM $H_2O_2$), and reducing (3 min, 5 mM DTT) conditions. Tukey-style box plots are presented, where boxes represent the 25th to 75th percentiles with the horizontal line indicating the median, while whiskers extend to the most extreme data points within 1.5 times the interquartile range (IQR) from the quartiles; points beyond this range are considered outliers. Statistical significance was evaluated by one-way ANOVA (Tukey's multiple comparisons test) with N > 60 independent cells examined, yielding a Golgi redox potential ($E_{GSH}$) of 144 ± 18 mV (pH = 6.2), calculated using the Nernst equation. **Right**: Representative fluorescent microscopy images acquired at excitation wavelengths of 405 nm (pseudo-colored in cyan) and 488 nm (pseudo-colored in green) with emission collected at 525/50 nm. Scale bar: 10 μm.

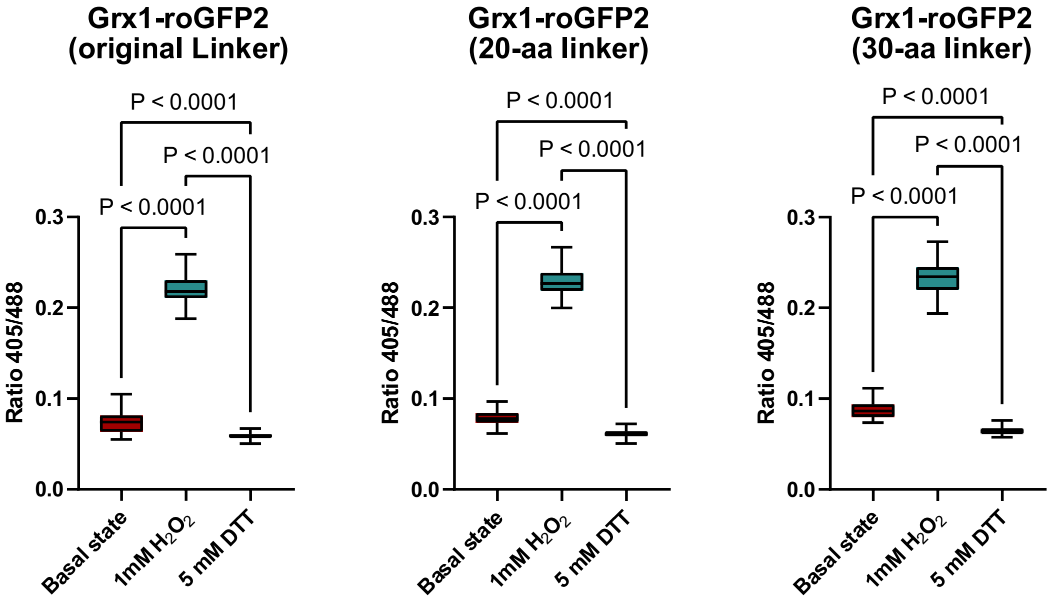

**Figure S2. Validation of the Redesigned Grx1-roGFP2 Biosensors.** Functional redox responsiveness of the original sensor and the redesigned sensors featuring 20- and 30-amino-acid linkers was assessed by the fluorescence ratio (405 nm / 488 nm) under basal (untreated), oxidizing (1 mM $H_2O_2$), and reducing (5 mM DTT) conditions. Tukey-style box plots are presented as boxes represent the 25th to 75th percentiles with the horizontal line indicating the median, while whiskers extend to the most extreme data points within 1.5 times the interquartile range (IQR) from the quartiles; any points beyond this range are considered outliers. Statistical significance was evaluated by one-way ANOVA (Šídák's multiple comparisons test) with N > 65 independent cells examined. The box plots demonstrate that all variants exhibit reversible shifts in the fluorescence ratio, confirming that the redesigned Grx1-roGFP2 constructs maintain robust redox sensitivity comparable to the original.

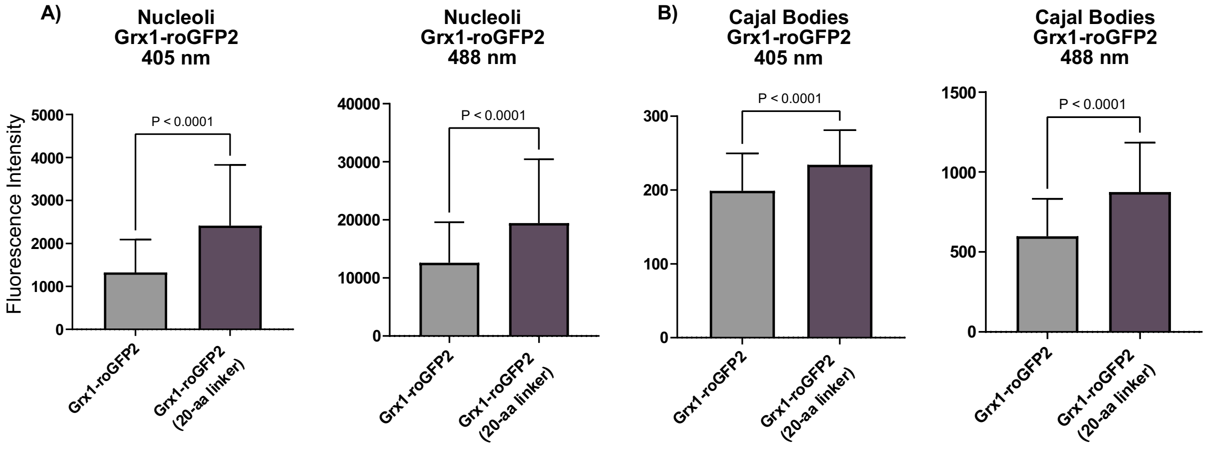


**Figure S3.** Quantitative bar charts comparing the mean fluorescence intensities (± SD) for the original Grx1-roGFP2 sensor and miniGrx1-roGFP2, as measured in the 405 nm and 488 nm channels for the **(A)** nucleoli and **(B)** Cajal bodies. The statistical significance was determined using an unpaired t-test.

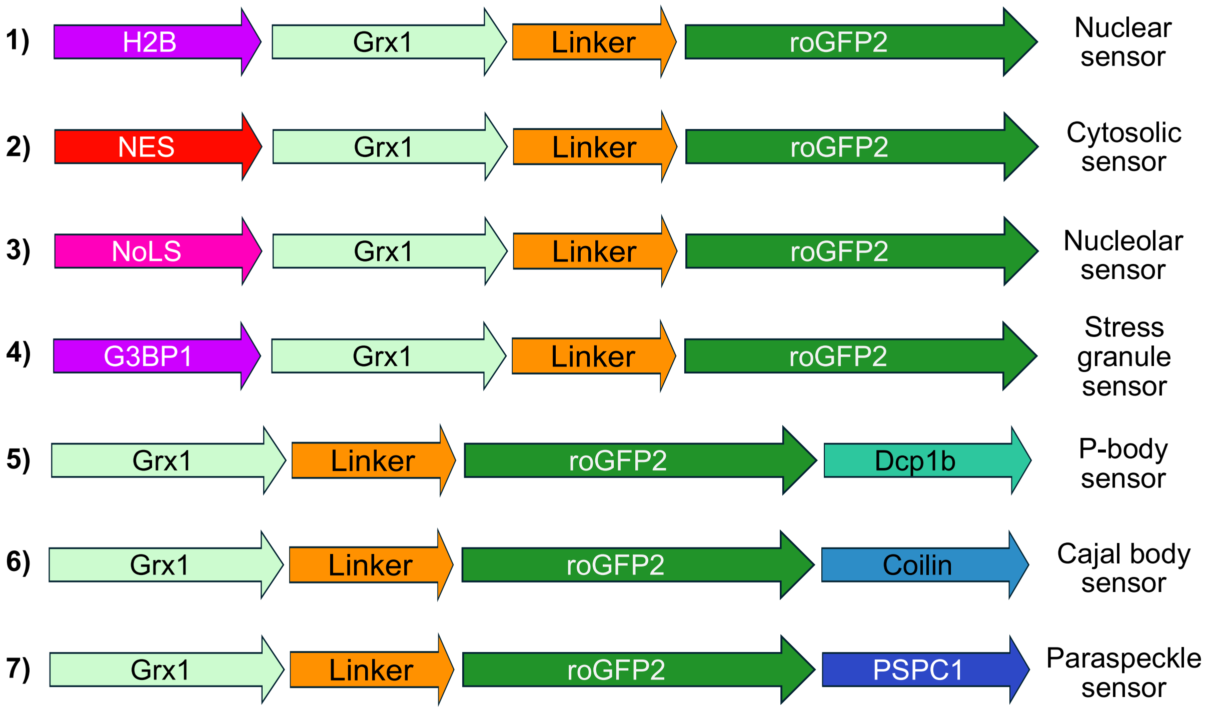


**Figure S4. Schematic representation of plasmid constructs targeting the miniGrx1-roGFP2 sensor to distinct subcellular compartments.** The constructs include: (1) a nuclear sensor, (2) a cytosolic sensor, (3) a nucleolar sensor, (4) a stress granule sensor, (5) a P-body sensor, (6) a Cajal body sensor, and (7) a paraspeckle sensor. Each construct is engineered with specific localization signals to direct the sensor to the corresponding compartment.

### 2. Supplementary Tables

**Table S1. $E_{GSH}$ Values Across Subcellular Compartments and MLOs.** $E_{GSH}$ values (in mV) are shown for stress granules, processing bodies, nucleoli, nuclear paraspeckles, and Cajal bodies, along with the corresponding surrounding compartments (cytosol for stress granules and processing bodies; nucleus for nucleoli, paraspeckles and Cajal bodies). Data were calculated using the Nernst equation. The standard deviations (SD) represent uncertainties derived from propagating the experimental errors through the Nernst equation.

<table>
<tr><th></th><th>Region</th><th>$E_{GSH}$ (mV)</th><th>Surrounding Compartment</th><th>$E_{GSH}$ (mV)</th></tr>
<tr><td rowspan="5">Replicate 2</td><td>Stress Granules</td><td>-282 ± 25</td><td rowspan="2">Cytosol</td><td rowspan="2">-304 ± 15</td></tr>
<tr><td>Processing Bodies</td><td>-287 ± 19</td></tr>
<tr><td>Nucleoli</td><td>-290 ± 16</td><td rowspan="3">Nucleus</td><td rowspan="3">-292 ± 12</td></tr>
<tr><td>Paraspeckles</td><td>-300 ± 12</td></tr>
<tr><td>Cajal Bodies</td><td>-290 ± 10</td></tr>
<tr><td rowspan="5">Replicate 3</td><td>Stress Granules</td><td>-293 ± 30</td><td rowspan="2">Cytosol</td><td rowspan="2">-295 ± 30</td></tr>
<tr><td>Processing Bodies</td><td>-287 ± 30</td></tr>
<tr><td>Nucleoli</td><td>-302 ± 15</td><td rowspan="3">Nucleus</td><td rowspan="3">-310 ± 11</td></tr>
<tr><td>Paraspeckles</td><td>-302 ± 15</td></tr>
<tr><td>Cajal Bodies</td><td>-296 ± 17</td></tr>
</table>

**Table S2.** Plasmid sources with the plasmid name, the source, vector precursor, and the insert precursor.

| Plasmid | Source | Vector precursor | Insert precursor |
|---|---|---|---|
| H2B-Grx1-roGFP2 | Gibson assembly | H2B-HT-mGold | GALT1-Grx1-roGFP2 |
| NES-Grx1-roGFP2 | Site-directed Mutagenesis | H2B-Grx1-roGFP2 | |
| H2B-Grx1-d-roGFP2 | Gibson assembly | H2B-Grx1-roGFP2 | H2B-Grx1-roGFP2 |
| Nols-Grx1-roGFP2 | Site-directed Mutagenesis | H2B-Grx1-roGFP2 | |
| G3BP1-Grx1-roGFP2 | Gibson assembly | H2B-Grx1-roGFP2 | G3BP1-iRFP[1] |
| Grx1-roGFP2 | Site-directed Mutagenesis | H2B-Grx1-roGFP2 | |
| Grx1-roGFP2-Dcp1b | Gibson assembly | Grx1-roGFP2 | GFP-Dcp1b[2] |
| Grx1-roGFP2-Coilin | Gibson assembly | Grx1-roGFP2 | EGFP-Coilin[3] |
| Grx1-roGFP2-PSPC1 | Gibson assembly | Grx1-roGFP2 | pCDNA3-HA-PSPC1[4] |
| Galt1-Grx1-20aaAlign-roGFP2 | Gibson assembly | Nols-Grx1-20aaAlign-roGFP2 | Galt1-HT7-mGold[5] |

**Table S3**. Primers for plasmid generation.

| Plasmid | Vector forward | Vector reverse | Insert forward | Insert reverse |
|---|---|---|---|---|
| H2B-Grx1-roGFP2 | gtacaagtaagagagctaagcggccgcgact | tgagccatgctgccggtcgactgctta | cggcagcatggctcaagagtttgtg | gcttagctctcttacttgtacagctcgtccatgc |
| NES-Grx1-roGFP2 | aactggaactggatgaaggaggacagtcgaccggcagcatg | cttccagcttcttttgcagagccatggtggcggtaccgtc | | |
| H2B-Grx1-d-roGFP2 | agcacctcctccgccctgagcaaag | acgttgtgggagttgtagttgtactccagcttgt | actacaactcccacaacgtctatatcatggccgac | agggcggaggaggtgctcaggtag |
| NoLs-Grx1-roGFP2 | caccgccgccgccgccgccgccgccagtcgaccggcag | caccggcaccttgcgcttcttctttggcatggtggcgg | | |
| G3BP1-Grx1-roGFP2 | acggcagcagtcgaccggcagca | tctccatcaccatggtggcggtaccgtcga | cgccaccatggtgatggagaagc | tcgactgctgccgtggcgcaa |
| Grx1-roGFP2 | atggctcaagagtttgtgaac | ggtggcggtaccgtc | | |
| Grx1-roGFP2-Dcp1b | gactatgtgagagagctaagcggccgcgact | ctgagtccggacttgtacagctcgtccatgc | cgagctgtacaagtccggactcagatctcg | gcttagctctctcacatagtctttttcatggctgct |
| Grx1-roGFP2-Coilin | gctgtacaagtactcagatccaccaag | gctctcttaggcaggttctgtacttgatgtgt | aacctgcctaagagagctaagcggc | gatctgagtacttgtacagctcgtccatgccg |
| Grx1-roGFP2-PSPC1 | gctgtacaagatgatgttaagaggaaacctgaagc | ccgcttagctctcttaatatctacgacgcttattaggg | gtcgtagatattaagagagctaagcggc | cttaacatcatcttgtacagctcgtccatgcc |
| Galt1-Grx1-20aaAlign-roGFP2 | gctgcagcagtcgaccggcagcat | cgaagcctcatggtggcggtaccgtc | caccatgaggcttcgggagcc | gtcgactgctgcagcggtgtggaga |

**Table S4.** Microscope settings for imaging channels.

| Channel | λ excitation | Emission filter | Laser intensity |
|---|---|---|---|
| roGFP2 blue | 405 nm | 525/50 | 1.65 mW |
| roGFP green | 488 nm | 525/50 | 1.24 mW |

**Table S5. Statistical Analysis for Figure 1B and Figure 1C.** Data for **Figure 1B** represent the one-way ANOVA (Tukey's multiple comparisons test) analysis of fluorescence ratio measurements (405 nm/488 nm) for the original and the 20-aa linker redesigned sensor under basal conditions, oxidizing conditions (1 mM $H_2O_2$), and reducing conditions (5 mM DTT). Data for **Figure 1C** compare the fluorescence intensities in the nucleus (basal state) among the original sensor, the 20-aa linker redesigned sensor, and the 30-aa linker redesigned sensor, as measured in the 405 nm and 488 nm channels. For each comparison, the mean difference, 95% confidence interval, and adjusted p-value are provided, indicating the statistical significance of the observed differences.

| Tukey's multiple comparisons test | | | Mean Diff. | 95.00% CI of diff. | Below threshold? | Summary | Adjusted P Value |
|---|---|---|---|---|---|---|---|
| Figure 1B | Original Linker | Basal state vs. 1mM $H_2O_2$ | -0.1481 | -0.1535 to -0.1427 | Yes | **** | <0.0001 |
| | | Basal state vs. 5 mM DTT | 0.01964 | 0.01434 to 0.02495 | Yes | **** | <0.0001 |
| | | 1mM $H_2O_2$ vs. 5 mM DTT | 0.1677 | 0.1625 to 0.1729 | Yes | **** | <0.0001 |
| | 20-aa Linker | Basal state vs. 1mM $H_2O_2$ | -0.1468 | -0.1508 to -0.1428 | Yes | **** | <0.0001 |
| | | Basal state vs. 5 mM DTT | 0.01511 | 0.01091 to 0.01931 | Yes | **** | <0.0001 |
| | | 1mM $H_2O_2$ vs. 5 mM DTT | 0.1619 | 0.1578 to 0.1661 | Yes | **** | <0.0001 |
| Figure 1C | 488 nm | Grx1-roGFP2 vs. Grx1-roGFP2 (20-aa linker) | -6495 | -9151 to -3839 | Yes | **** | <0.0001 |
| | | Grx1-roGFP2 vs. Grx1-roGFP2 (30-aa linker) | -4583 | -7102 to -2063 | Yes | **** | <0.0001 |
| | | Grx1-roGFP2 (20-aa linker) vs. Grx1-roGFP2 (30-aa linker) | 1912 | -897.1 to 4721 | No | ns | 0.2454 |
| | 405 nm | Grx1-roGFP2 vs. Grx1-roGFP2 (20-aa linker) | -578 | -812.8 to -343.3 | Yes | **** | <0.0001 |
| | | Grx1-roGFP2 vs. Grx1-roGFP2 (30-aa linker) | -495.7 | -718.4 to -273.1 | Yes | **** | <0.0001 |
| | | Grx1-roGFP2 (20-aa linker) vs. Grx1-roGFP2 (30-aa linker) | 82.32 | -166.0 to 330.6 | No | ns | 0.7145 |

**Table S6. Statistical Analysis for Figure 2B**. One-way ANOVA with Tukey's multiple comparisons test was performed on the fluorescence ratio data (405 nm/488 nm) measured under basal, oxidizing (1 mM $H_2O_2$), and reducing (5 mM DTT) conditions across various subcellular compartments. The table lists the mean differences, 95% confidence intervals, and adjusted p-values for comparisons within each compartment (nucleus, nucleoli, Cajal bodies, paraspeckles, cytosol, stress granules, and p-bodies), highlighting significant differences in redox responses.

| | | | | | | |
|---|---|---|---|---|---|---|
| Nucleus | Basal state vs. 1mM $H_2O_2$ | -0.4443 | -0.4562 to -0.4324 | Yes | **** | <0.0001 |
| | Basal state vs. 5 mM DTT | 0.03691 | 0.02603 to 0.04778 | Yes | **** | <0.0001 |
| | 1mM $H_2O_2$ vs. 5 mM DTT | 0.4812 | 0.4700 to 0.4925 | Yes | **** | <0.0001 |
| Nucleoli | Basal state vs. 1mM $H_2O_2$ | -0.4286 | -0.4377 to -0.4195 | Yes | **** | <0.0001 |
| | Basal state vs. 5 mM DTT | 0.03947 | 0.03036 to 0.04858 | Yes | **** | <0.0001 |
| | 1mM $H_2O_2$ vs. 5 mM DTT | 0.468 | 0.4589 to 0.4772 | Yes | **** | <0.0001 |
| Cajal bodies | Basal state vs. 1mM $H_2O_2$ | -0.474 | -0.4977 to -0.4504 | Yes | **** | <0.0001 |
| | Basal state vs. 5 mM DTT | 0.03539 | 0.01174 to 0.05904 | Yes | ** | 0.0015 |
| | 1mM $H_2O_2$ vs. 5 mM DTT | 0.5094 | 0.4843 to 0.5346 | Yes | **** | <0.0001 |
| Paraspeckles | Basal state vs. 1mM $H_2O_2$ | -0.48 | -0.4979 to -0.4622 | Yes | **** | <0.0001 |
| | Basal state vs. 5 mM DTT | 0.06375 | 0.04446 to 0.08304 | Yes | **** | <0.0001 |
| | 1mM $H_2O_2$ vs. 5 mM DTT | 0.5438 | 0.5238 to 0.5638 | Yes | **** | <0.0001 |
| Cytosol | Basal state vs. 1mM $H_2O_2$ | -0.4542 | -0.4659 to -0.4424 | Yes | **** | <0.0001 |
| | Basal state vs. 5 mM DTT | 0.03923 | 0.02746 to 0.05100 | Yes | **** | <0.0001 |
| | 1mM $H_2O_2$ vs. 5 mM DTT | 0.4934 | 0.4813 to 0.5054 | Yes | **** | <0.0001 |
| Stress Granules | Basal state vs. 1mM $H_2O_2$ | -0.4765 | -0.4897 to -0.4634 | Yes | **** | <0.0001 |
| | Basal state vs. 5 mM DTT | 0.06471 | 0.05181 to 0.07760 | Yes | **** | <0.0001 |
| | 1mM $H_2O_2$ vs. 5 mM DTT | 0.5412 | 0.5285 to 0.5539 | Yes | **** | <0.0001 |
| P-bodies | Basal state vs. 1mM $H_2O_2$ | -0.2959 | -0.3048 to -0.2870 | Yes | **** | <0.0001 |
| | Basal state vs. 5 mM DTT | 0.08763 | 0.07885 to 0.09641 | Yes | **** | <0.0001 |
| | 1mM $H_2O_2$ vs. 5 mM DTT | 0.3835 | 0.3732 to 0.3939 | Yes | **** | <0.0001 |

### 3. Experimental section

*Gibson assembly*

Simulation of the Gibson assembly was conducted using SnapGene (v4.1.9). Initially, primers for amplification were designed within SnapGene and subsequently adjusted by hand to reduce issues such as hairpin loops, self-dimer formation, sequence repeats, and imbalanced GC content. Details regarding the precursor plasmids and primer sequences are provided in Tables S2 and S3. All reagents and general protocols followed those included in the Gibson Assembly Cloning Kit (New England Biolabs). Vector and insert fragments were generated by PCR linearization using Phusion High Fidelity PCR Master Mix, with any residual template DNA eliminated via DpnI digestion. Following verification by gel electrophoresis, the PCR products were purified using the QIAquick PCR Purification Kit (Qiagen). The purified backbone and insert fragments were then ligated and introduced into NEB 5α competent cells through a heat shock transformation, following the vendor's standard procedures. Amplification of the resulting plasmids was carried out by incubating LB cultures containing the appropriate antibiotics at 37°C overnight. Finally, plasmid extraction and purification were completed using either the Qiagen Plasmid Mini kit or the Qiagen Plasmid Plus Midi kit, and the sequence of the gene of interest was confirmed via Sanger sequencing performed by Microsynth AG.

*Site-directed mutagenesis*

Site-directed mutagenesis was performed by first designing primers using the NEBaseChanger online tool. The precursor plasmids and corresponding primer sequences are detailed in Tables S2 and S3. Using the Site-Directed Mutagenesis Kit (NEB) with Phusion High Fidelity PCR Master Mix, the plasmids were amplified, followed by enzymatic digestion and ligation to introduce the desired mutations. The resulting constructs were then transformed into NEB 5α competent cells via heat shock following the vendor's standard protocol. Finally, the gene of interest was verified by Sanger sequencing performed by Microsynth AG.

*Cell culture*

HeLa cells were maintained in Dulbecco's Modified Eagle Medium (DMEM) that was enriched with 10% fetal bovine serum (FBS) and a mixture of penicillin (100 U $mL^{-1}$), streptomycin (100 µg $mL^{-1}$), and fungizone (0.25 µg $mL^{-1}$) at 37 °C under a 5% $CO_2$ atmosphere. For snapshot imaging, cells were seeded at a density of 15,000–20,000 per well in an 8-well Ibidi chambered cover glass 2–3 days before imaging, whereas for timelapse imaging, the seeding density was adjusted to 5,000–10,000 cells per well. Transfections were carried out using jetPRIME with plasmid DNA, following the supplier's protocol, one day prior to imaging. Before acquiring images, the growth medium was removed and cells were washed twice with PBS, then imaged in FluoroBrite DMEM for standard conditions. To induce stress granule formation, cells were incubated at 43 °C for 30 minutes prior to imaging. For timelapse experiments, imaging was performed in fresh FluoroBrite DMEM supplemented with 4 mM glutamine, 1 mM sodium pyruvate, 10% FBS, and the same antibiotics as described above.

*Confocal microscopy*

Confocal imaging was carried out on a dual-camera Nikon W1 spinning disc microscope fitted with an sCMOS camera (Photometrics). Brightfield images were obtained using a white LED, while fluorescence was excited with 405 nm and 488 nm lasers, with appropriate filter cubes configuring the light path. High-resolution images were captured using a CFI Plan Apo VC water immersion objective (60×, NA = 1.2), whereas timelapse experiments utilized a 20× air objective. Imaging was performed sequentially for each channel, and the system was controlled via NIS Elements software. All imaging sessions were conducted at 37 °C in a 5% $CO_2$ environment, and subsequent image analysis was performed using Fiji/ImageJ.

*Image analysis*

The image analysis was performed based on an extended version of the ratiometric image analysis tool developed in Emmert *et al.* (2023).[7] This tool uses thresholding to differentiate between image background and objects in each image, where each image has two channels depending on the application. After identifying the objects in the images, the tool then calculates the ratio between the channels for each pixel of the identified objects, which can then be used to calculate the GSH redox potential.

In this project, we adapted the tool by automatizing the timelapse analysis, adding a cell separation algorithm, the ability to identify secondary objects, and the masking procedure as follows:

- Timelapse analysis: The script used to calculate the ratiometric values are the same for timelapses and snapshots. The tool automatically sorts the images and analyzes them in order across timeframes.
- Cell separation: The script is able to separate objects that are touching each other. For this, we follow the algorithm used in CellProfiler (Stirling *et al.,* 2021).[8] Concretely, the script first calculates for each object the distance of all pixels to the nearest background pixel. Second, it identifies the local maxima within each object using a disc structuring element whose size is defined based on the average object size in the image. Finally, after identifying the possible local maxima within an object given the size of the structuring element, the tool uses the watershed algorithm to separate the object.
- Secondary objects: The script has the option to identify secondary objects within an already identified object. Here, we use a simple thresholding approach where the pixels with an intensity in the upper 20th percentile of each primary object are marked as secondary objects. This approach was implemented to separate nucleoli from nuclei.
- Masking procedure: To avoid having objects that are only present in one channel but not the other (e.g., dust) we use a composite mask to identify objects, i.e., we identify

objects in both channels, and only keep the pixels that are identified as objects in both channels. After identifying the object, we then compute the background value as the average of all values that are outside of the identified objects.

As our extended tool was not able to identify Cajal bodies and paraspeckles due to their small size, their identification and subsequent calculation of their ratio was done manually using Fiji/ImageJ. To this end, we manually marked the organelle, retrieved the fluorescence intensity from both channels within the marked area, subtracted the background, and finally calculated the ratio.
